# Simvastatin mediates inhibition of exosome synthesis, localization and secretion via multicomponent interventions

**DOI:** 10.1101/366211

**Authors:** Ankur Kulshreshtha, Swati Singh, Kritika Khanna, Anurag Agrawal, Balaram Ghosh

**Author notes:** **To whom all correspondence should be addressed**: **Balaram Ghosh**, Molecular Immunogenetics Laboratory, CSIR-Institute of Genomics and Integrative Biology, Mall Road, Delhi 110007, India, **Ankur Kulshreshtha**, Molecular medicine group, International Center for Genetic Engineering and Biotechnology, Aruna Asaf Ali Marg, Delhi 110 067, India.

## Abstract

Discovery of exosomes as modulator of cellular communication has added a new dimension to our understanding of biological processes. Exosomes influence the biological systems by mediating trans-communication across tissues and cells, which has important implication for health and disease. Identification of strategies for exosome modulation may pave the way towards better understanding of exosome biology and development of novel therapeutics. In absence of well-characterized modulators of exosome biogenesis, an alternative option is to target pathways generating important exosomal components. Cholesterol represents one such essential component required for exosomal biogenesis. We initiated this study to test the hypothesis that owing to its cholesterol lowering effect, simvastatin, a HMG CoA inhibitor, might be able to alter exosome formation and secretion. Using previously established protocols for detecting secreted exosomes in biological fluids, simvastatin was tested for its effect on exosome secretion under various in-vitro and in-vivo settings. Murine model of AAI was used for further validation of our findings. Utilizing aforementioned systems, we demonstrate exosome-lowering potential of simvastatin in various in-vivo and in-vitro models, of AAI and atherosclerosis. We believe that the knowledge acquired in this study holds potential for extension to other exosome dominated pathologies and model systems.

## Introduction

Exosomes are cell secreted membrane bound nano-vesicular structures that have been shown to modulates the function and phenotype of recipient cells, ^1^ via transfer of associated lipids, proteins, RNA, and DNA species. ^2,3^ Such contents vary with cell lineage and state, accounting for a wide range of reported effects. ^4^ For example, stem cell exosomes render protective effects, while ^5^ cancer cell derived exosomes promote metastasis. ^6,7^ Pro-inflammatory role for exosomes has also been demonstrated in other pathological conditions. ^8–10^ some studies have even proposed that strategies to reduce exosome secretion might have protective effects during inflammatory conditions. ^11,12^

Formation and secretion of exosomes is a complex biological process, detailed knowledge of which remains incomplete. Though recent studies have started identifying key proteins involved in this process, such as PI3K, Akt, eNOS, Alix, syndecan, syntenin, Rab-27a and Rab-27b, ^12–14^ their precise role in this complex process is still under investigation. Other than proteins, ceramide and calcium have also been reported to regulate exosome biogenesis and secretion respectively. ^15,16^ Despite rapidly emerging evidence for associative role of exosomal communication in inflammatory diseases, there has been little progress towards identification of drug-candidates that can inhibit exosome secretion. While experimental studies have used siRNAs against important proteins ^12^ and pharmacological inhibitors such as GW4869 ^16^, they still await approval for human use. Towards filling this lacuna, we reasoned if inhibition of cholesterol synthesis by statins could be a viable strategy for inhibiting exosome secretion, as cholesterol is most abundant component of exosomal membrane and statins represent safe and approved class of drugs for limiting cholesterol availability. This approach seemed plausible for three reasons: a) cholesterol is a necessary lipid precursor for formation of exosomal membranes, and could offer a better target than proteins such as various Rab family proteins, whose exosome-independent functionality has not been explored yet, b) although statins are mainly employed for cholesterol biosynthesis inhibition, they also have a number of poorly understood additional anti-inflammatory effects ^17^ and, c) repurposing of an existing drug for exosome reduction would be far more fast and efficient toward clinical application than discovering novel drug candidates.

Here, we investigated if simvastatin could reduce formation and/ or secretion of exosomes, and whether this could offer protection against exosome mediated pro-inflammatory response in experimental models of asthma and in-vitro model of atherosclerosis. Our current data supports a novel mode of action for simvastatin in inhibiting both exosome formation and secretion that explains some poorly understood aspects of anti-inflammatory effects of statins and can be further utilized in several exosome-mediated inflammatory conditions.

## Results

### Simvastatin reduces exosome secretion *in vitro*

Important role for cholesterol in formation of exosomes has been previously reported, ^16^ so we reasoned if limiting cholesterol availability in target cells could hinder exosome production. Literary evidences wherein cholesterol reduction has been shown to impair exocytosis of synaptic vesicles support this hypothesis. ^18^ To further test this hypothesis, we treated exosome-secreting cells with simvastatin, and measured effect on exosome secretion using a semi-quantitative fluorescent bead-based assay ^10^ and for exosome associated proteins using western-blotting. We also validated the reduction in cholesterol levels upon simvastatin treatment **(Figure S1 in the online supplement)**. Characterization of these particles as exosomes using EM, western blotting, DLS and density gradient centrifugation has already been described in an earlier report from our group, but we still validated the size and morphology using TEM **(Figure S2 in the online supplement)**. ^10^ Epithelial cells and monocytes treated with increasing concentration of simvastatin for a period of 24 hours exhibited a significant reduction in the level of secreted exosomes, as measured by the bead-based assay **A-B).** A significant reduction of about 40% was noted at the 0.3 μM dose of simvastatin, which corresponds to non-toxic maximal plasma concentration associated with simvastatin therapy in humans, ^19^ confirming the plausibility of this effect at usual clinical dosing. The lack of toxicity was also confirmed by MTT staining (data not shown) and visible inspection **(Figure S3 in the online supplement)**. The efficacy of bead-based assay was validated by confirming linearly increased detection of exosome-associated proteins, such as CD9/CD81 and Annexin-V, in cell-culture supernatants from increasing number of cells **(Figure S4 in the online supplement)**. These effects were confirmed further by measuring exosome-associated proteins, Alix, Tsg-101 and β-actin in pelleted exosome fraction of culture supernatant from lowest effective dose (0.3 μM) of simvastatin **(Figure 1C)**.

**Figure 1.**
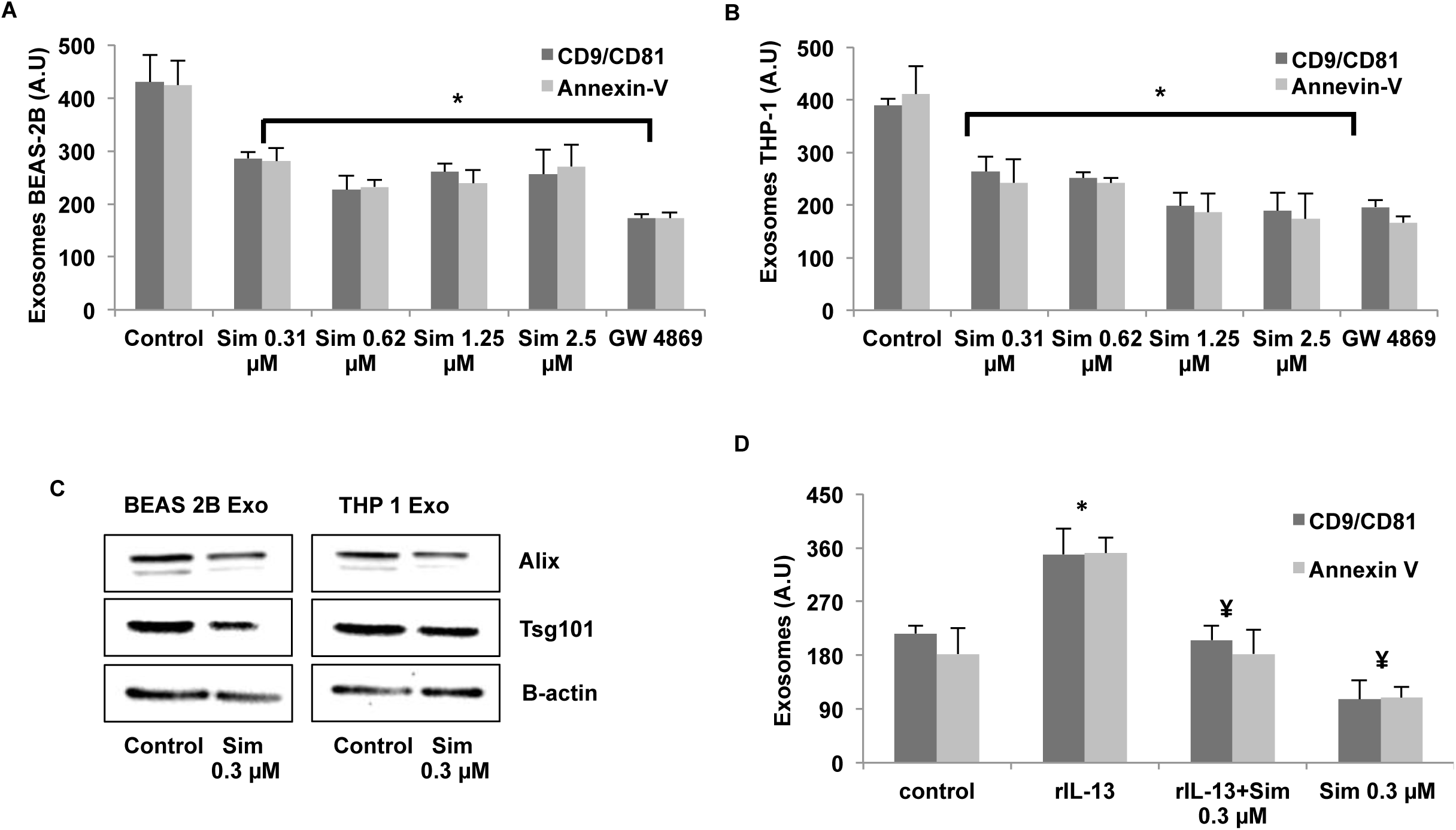
Simvastatin reduces exosomes secretion. **(A-B)**, Cells at concentration of 1×10^6/ ml were treated with indicated concentrations of simvastatin in 2 ml of media for a period of 24 hours, after which the culture supernatant was harvested and 1ml from it was used for measuring exosomes. Secreted exosome levels in culture supernatant from simvastatin treated epithelial cells **(A)** and THP-1 monocytes **(B)**, measured as in **Figure E1, A in the online repository. (C)**, Levels of exosome associated Alix, Tsg-101 and βactin in pelleted exosome fraction from supernatant of 10×10^6 simvastatin treated cells. **(D)** Effect of simvastatin treatment on exosome associated CD9/CD81 and Annexin V in cell culture supernatant from IL13 (25 ng/ml) and simvastatin treated epithelial cells. Data in **A, B** and **D** represent the mean±SE from three independent experiments. Data in **C** is representative image from one of the two independent experiments. (∗p<0.05 vs Control and ¥ p< .05 vs rIL13). Sim: Simvastatin.

In an earlier study, we had demonstrated that IL-13 treatment led to increased production of pro-inflammatory exosomes from airway epithelial cells. ^10^ We tested if simvastatin treatment could reverse this process as well, and observed that simvastatin treatment significantly reduced the levels of secreted exosomes from IL-13 treated epithelial cells **(Figure 1D)**, as measured by bead-based assay mentioned above. This corroborates well with previous observations wherein simvastatin has been shown to play a beneficial role in asthma ^20,21^ and we propose that inhibition of proinflammatory exosomes could be one of the mechanisms behind simvastatin’s protective effects.

### Simvastatin reduces the intracellular levels of exosome-associated proteins

Our initial data suggested that simvastatin treatment led to reduction in exosome secretion, possibly by conventional action of cholesterol reduction by statin **(Figure S1 in the online supplement)**, but the detailed mechanism behind it remain unclear. Few reports have identified eNOS and Alix axis in exosome biogenesis wherein Alix has been shown to positively regulate the exosome secretory process ^12,13^ and a negative regulatory role for eNOS. ^12^ We had previously demonstrated that simvastatin increases eNOS levels, both in cultured airway epithelial cells, as well as in *vivo*, ^22^ we further sought to determine if it was also affecting ALIX levels. We observed that simvastatin, but not another exosome inhibitor (GW4869), reduce the levels of Alix and CD-63 **(Figure 2A).** Surface levels of important exosome associated proteins CD63 and CD81 but not E-cadherin, were also dose-dependently reduced by simvastatin **(Figure 2B),** confirming the specificity and general reduction in these proteins by simvastatin treatment. The reduction in exosome-associated proteins like CD63 was additionally confirmed by immunofluorescence microscopy **(Figure 2C)**. Thus, reduction in levels of exosome synthesizing proteins may partly contribute to the simvastatin-mediated reduction in exosome secretion.

**Figure 2.**
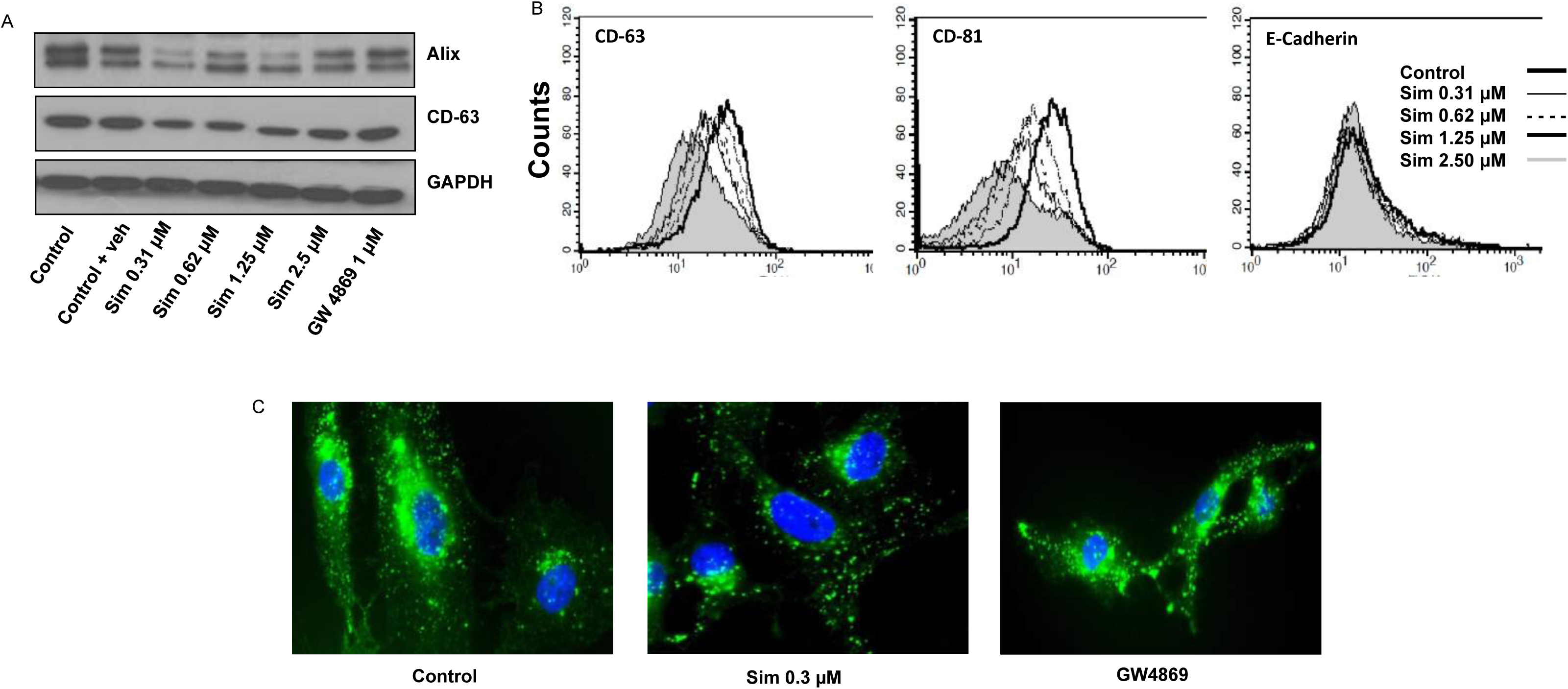
Simvastatin directly alters the level of various exosome associated proteins. **(A)** Western blots for Alix and CD63 levels in total cell protein with different doses of simvastatin. **(B)** Cell surface levels of CD63 and CD81 were measured by flow cytometry after treatment with various doses of simvastatin, E-cadherin was used as control surface marker. **(C)** Immunocytochemistry for CD63 on cells treated with indicated concentration of simvastatin. Sim: Simvastatin.

### The effect of simvastatin on exosome production and inflammation is only partially related to cholesterol reduction

We (and others) have previously reported elevated levels of pro-inflammatory exosomes in asthmatic lung as well as in BALF from mice of experimental model of allergic airway inflammation (AAI). Pharmacological inhibition of these exosomes was found to provide protective effect in experimental asthma. ^10^ To determine whether simvastatin treatment would inhibit exosome secretion and in turn attenuate AAI, we used a well-established mouse model of AAI **(Figure S5 in the online supplement)**. Further, to determine whether the effect of simvastatin was due to inhibition of the mevalonate formation step of cholesterol biosynthesis, or an independent pleiotropic effect, we additionally administered excess mevalonic acid ^23^ to a group of simvastatin-treated mice with AAI. Increased exosome content in BAL fluid in AAI was fully reversed by simvastatin treatment **(Figure 3A)**. However, a large residual effect of simvastatin was seen even after mevalonate supplementation. This reduction in exosome content was found to be associated with protective effect on other asthmatic features as well, such as inflammation, mucin granule production, AHR, cell-count and serum IgE **(Figure 3B-G)**. Also, in ova-challenged mice, simvastatin significantly decreased the levels of IL-4, IL-13 **(Figure 3H)**, IL-5 and IL-10 **(Figure S6, A-B in the online supplement)**. Mevalonate co-treatment, however, did not reverse the inhibitory effect of simvastatin on these cytokines. Levels of IFN-γ, an important Th1 cytokine, was unperturbed by any of the treatments **(Figure S6, C in the online supplement)**. Thus the effect of simvastatin on exosomes and AAI in mouse model could only be partly explained by reduced cholesterol synthesis and may relate to other pleiotropic effects as suggested previously. As animal models are complex systems, often involving interaction of several components leading to less clean readouts, we further validated the exosome inhibitory potential of simvastatin in a simpler in-vitro interaction system mimicking pro-atherogenic exosomal interaction between monocytes and endothelial cells.

**Figure 3.**
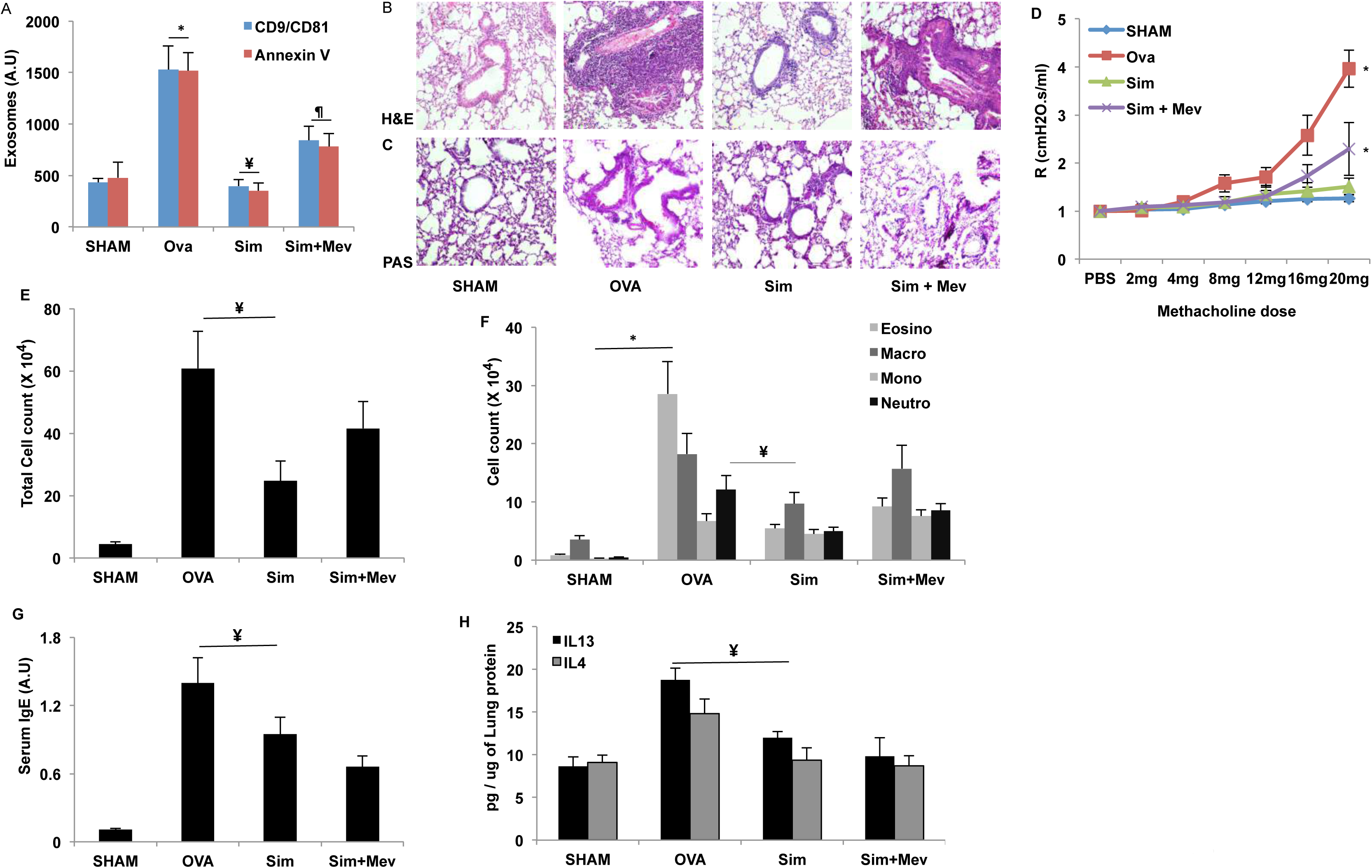
Effect of simvastatin and mevalonate cotreatment on inflammatory parameters. **(A)** Secreted exosome levels in BAL supernatant of mice from indicated groups. **(B-C)** Lung sections stained with hematoxylin and eosin **(H&E, B)** showing leukocyte infiltration, periodic acid–Schiff **(PAS, C)** for collagen deposition. **(D)** Airway resistance with increasing concentrations of methacholine 12h after the last challenge. **(E-F)** Effect of indicated treatments on total leukocyte count **(E)** and differential leukocyte count enumerated by morphological criteria **(F)**. **(G)** Ova specific serum IgE levels measured by ELISA. **(H)** Cytokines IL-13 and IL-4 measured in pulmonary homogenate. Stains in **(B, C)** shown at 20X magnification. **Br**, Bronchus. Results **(A, D, E, F, G, H)** are the mean ± SE for each group from two experiments with 4-6 mice in each group, (**^∗^**, p<0.05 vs SHAM and ¥ p<0.05 vs OVA.), Sim: Simvastatin (40 μmg/kg/dose), Mev: Mevalonate (20 mg/kg/dose).

### Simvastatin mediated reduction in monocytic exosomes renders a protective effect in an *in vitro* model of atherosclerosis

Atherosclerotic plaque formation is a process whereby deposition of excess lipid and cholesterol in coronary artery leads to narrowing of blood vessels, thereby causing a reduction in blood flow to heart, resulting in heart failure. Atherosclerotic lesions are usually characterized by increased endothelial migration. In a study exploring this phenomenon, ^24^ authors implicated the role of exosomes (referred to as microvesicles in this paper) secreted by plaque-associated monocytes in endothelial migration. Microvesicle (MV) associated mir-150 was identified as the key driver of this process. ^24^ Since simvastatin has long been prescribed to patients of cardiovascular disorders, we wondered if one of the mechanisms by which it renders its protective effects could be by inhibiting microvesicle secretion from accumulated monocytes at plaque surface. For testing this hypothesis, we adopted the model previously described, ^24^ wherein monocytic microvesicles were shown to promote endothelial migration, and in turn atherosclerosis. These microvesicles contained several micro-RNA species including mir-150, mir-16 and mir-181a, however the pro-atherogenic nature of these vesicles was attributed majorly to mir-150, which caused reduction of c-myb in nearby endothelial cells, hence promoting their migration from the site of plaque formation.

Simvastatin treatment of monocytic cell line, THP-1, led to reduction in exosome secretion **(Figure 1C)**, and a consequent reduction in the levels of secreted mir-150 **(Figure 4A)**. mir-16 and mir-181b were used as positive controls for exosomes-associated micro-RNA content. Simvastatin treatment however did not significantly alter the intracellular levels of any of these miRNAs **(Figure 4B).** Incubation of THP-1 derived DIO labeled MVs with HUVECs led to rapid uptake of these vesicles by HUVECs **(Figure 4C)**, resulting in increased levels of mir-150 **(Figure 4D)**. Treatment of monocytes with simvastatin led to reduction in number of secreted microvesicles, and hence reduction in microvesicle-acquired mir-150 in HUVECs. mir-150 has been demonstrated to promote endothelial migration ^24^ and we also observed similar phenomenon in HUVECs treated with THP-1 derived microvesicle, in presence or absence of serum as a chemoattractant **(Figure 4E and Figure S7 in the online supplement)**. Treatment of THP-1 with simvastatin before MV isolation significantly reduced migration of HUVECs, exhibiting an atheroprotective phenotype **(Figure 4E2)**. Our results thus suggest that inhibition of monytic exosomes could be one of the alternate mechanism by which simvastatin renders a protective role in atherosclerosis.

**Figure 4.**
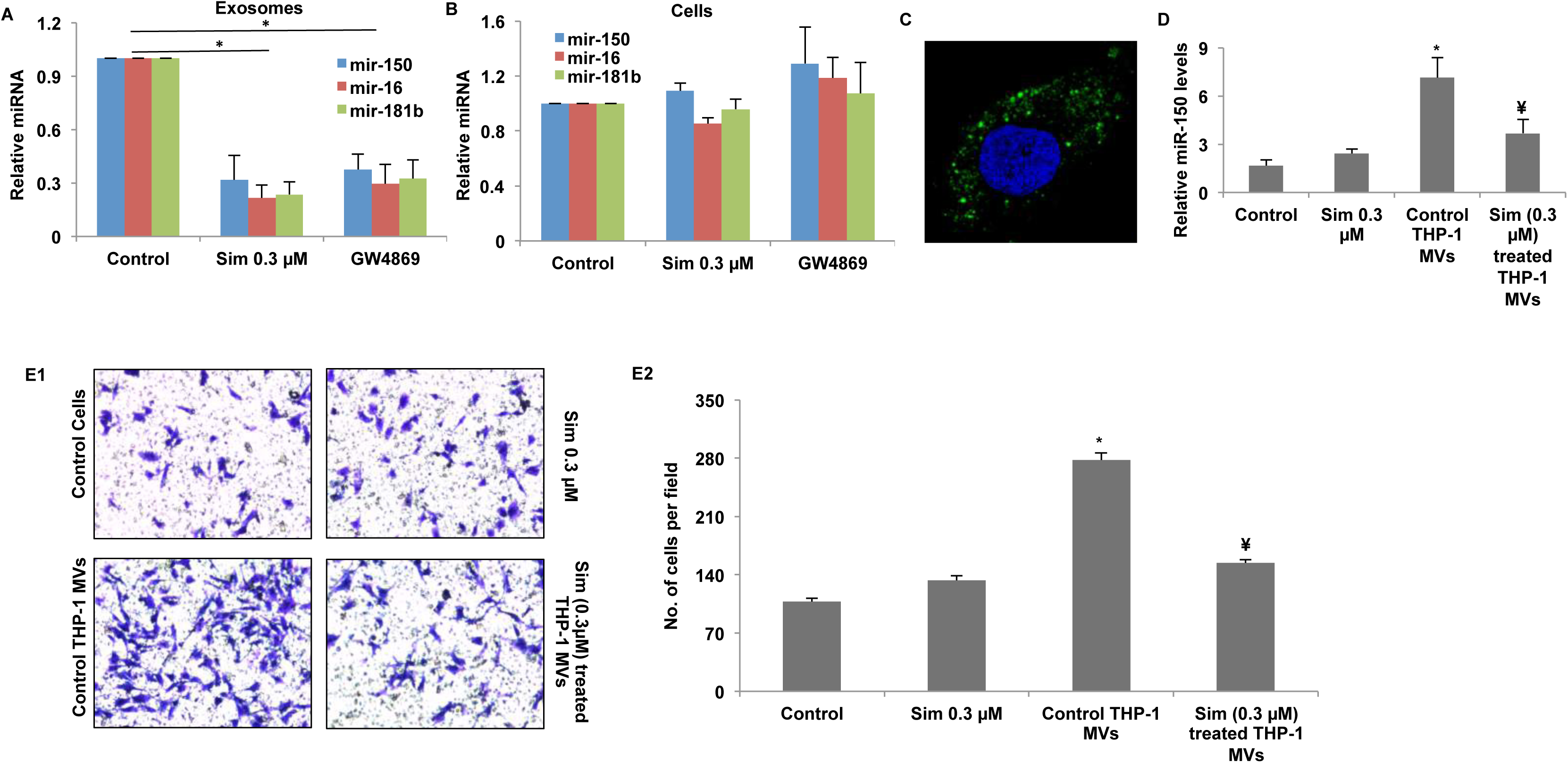
Simvastatin reduces exosome production from monocytes and attenuates exosome-enclosed mir-150 mediated endothelial cell migration. 1x10^6^ of THP-1 cells were seeded and treated with 0.3μM of simvastatin and 2μM GW4869 for a period of 24 hours. Cell pellet and supernatant was harvested and used separately for RNA isolation. Presence of indicated micro-RNAs was determined using qRT-PCR. Simvastatin mediated reduction of exosomes secretion from THP-1 monocytes results in lower levels of secretory miRNAs **(A)** but not intracellular miRNAs **(B). (C)** Uptake of DIO-labeled THP-1 derived exosomes (10 μg/mL) by HUVECs. **(D)** Relative mir-150 levels in HUVECs with indicated treatment **(E),** Simvastatin mediated reduction in exosome secretion by THP-1 monocytes results in lower mir-150 levels in HUVECs incubated with exosomes from simvastatin treated THP-1 in comparison to exosomes from same number of untreated THP-1 control cells, and consequent reduction in migration of endothelial cells. (^∗^p<0.05 vs Control in **A**, **D** and **E2. ¥** p<0.05 **vs Control THP-1 MVs in D and E2). Sim**: Simvastatin, **Control:** Control HUVEC

Once the exosome inhibitory role of simvastatin was established using various model systems, we furthered our study to investigate the putative underlying molecular mechanism.

### Simvastatin treatment alters MVB trafficking and results in their accumulation near the plasma membrane

We found notable reduction in cellular CD-63 levels upon simvastatin treatment in *in vitro* systems **(Figure 2C)**, that led us to test if this observation extends to *in vivo* conditions as well. For this purpose, lung tissue sections from our mouse model of AAI were stained for CD-63. Under these conditions, lungs are known to have elevated exosome-associated proteins in epithelial cells and macrophages. ^10^ While inspecting lung-tissue sections of simvastatin treated mice, we observed an interesting phenomenon, wherein simvastatin treatment led to accumulation of CD-63 positive compartments near the plasma membrane in epithelial cells **(Figure 5A)** as well as in macrophages **(Figure 5B).**

**Figure 5.**
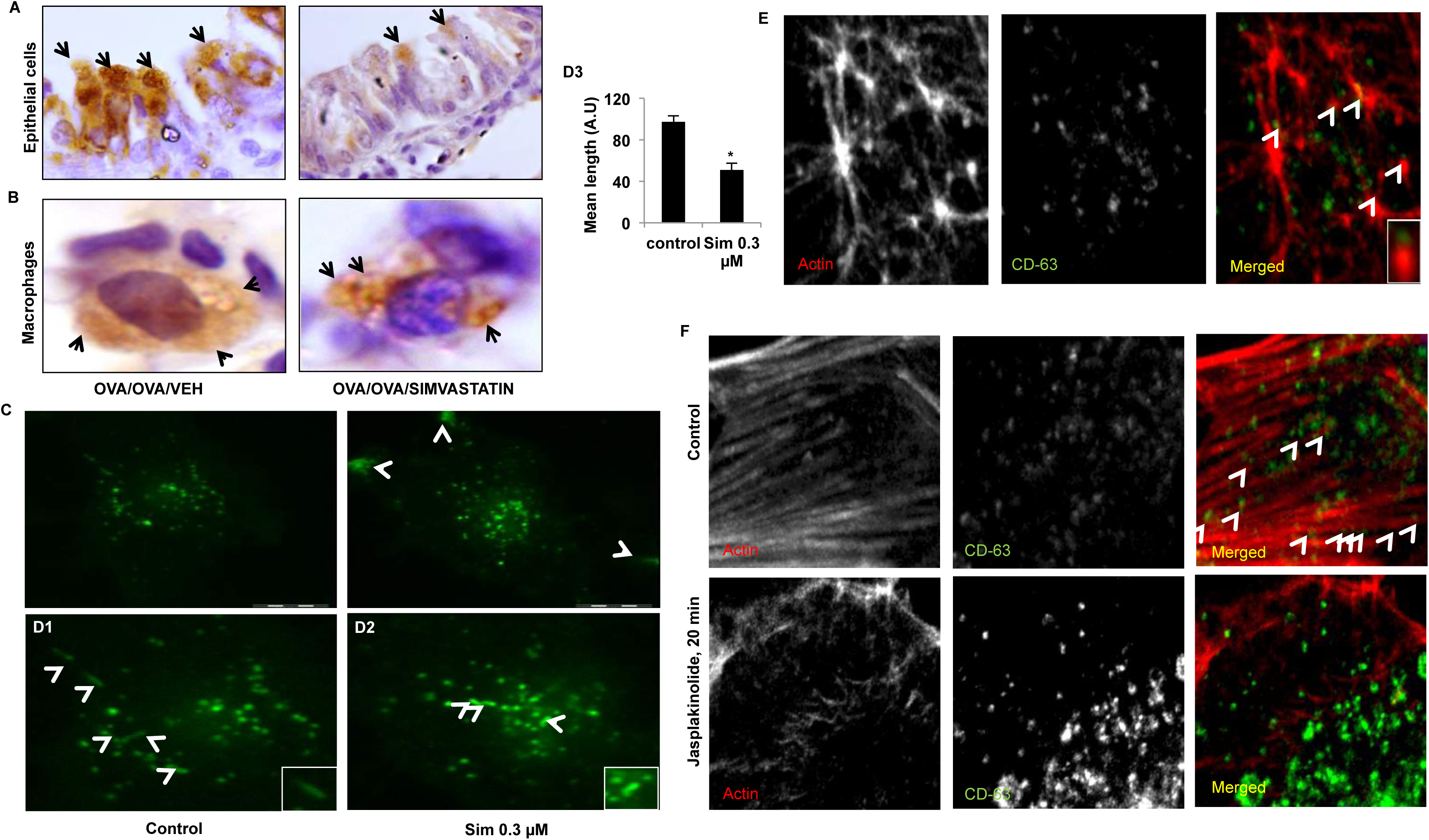
Simvastatin alters localization of CD63-positive compartments in cells. Representative Immunohistochemistry images for CD63 from lung tissue sections of OVA and OVA/Simvastatin treated mice in **(A)** epithelial cells and macrophages **(B)**. **(C),** Representative confocal images of CD-63 levels and localization post treatment with simvastatin. Arrows indicates CD63 localization pattern. **(D)** Representative images showing CD63-EGFP distribution and its association with linear beeline like structures **(D1-D2, inset)** in control and simvastatin treated cells in subplasmalemmal region, detected by TIRF microscopy and quantification of relative length **(D3). (E)** Co-localization of CD63 with actin nucleation sites as visualized by confocal microscopy. **(F)** Localization of CD63 with actin in absence and presence of actin polymerization enhancer Jasplakinolide (100 nM). Images in **(E, F)** shown at 63X while **(A, B, C, D)** shown at 100 X magnification. (∗p<0.05 vs Control in **Figure D3**). Sim: Simvastatin.

Since we observed an accumulation of CD63 positive compartments near the plasma membrane, we sought to investigate the fate of these compartments by using TIRF microscopy of CD63-EGFP transfected cells. While we found uniform distribution of MVBs and normal movement pattern in control cells, simvastatin treated cellsx had accumulation of MVBs near plasma membrane and restricted movement of other CD63 positive compartments, mostly towards the center of the cell **(Figure 5C, movie 1 and movie 2 in the online supplement)**.

Interestingly, while analyzing the TIRF data we found several CD63 positive compartments aligning with each other in a beeline pattern during their movement, suggesting them to be associated with well-defined cytoskeletal structures. These trails were shortened in simvastatin treated cells but had higher fluorescent intensity **(inset Figure 5D1-D2 and Figure 5D3)**.

Actins tracks have recently been implicated in movement of Rab11 and CCL2 containing vesicles, which initiate actin nucleation and elongation for their movement in a directed fashion. ^25^ We observed association of the CD63 positive compartments with actin nucleation sites **(Figure 5E)**. In light of these observations, we speculated that these CD-63 containing MVBs might be utilizing actin machinery for their movement. Closer examination of CD63 and actin inside cells revealed that the CD63 positive signals were indeed lying along the actin filaments and enhancing polymerization of actin structures by Jasplakinolide led to accumulation of CD-63 positive vesicles **(Figure 5F)**. Thus, we discovered that CD63 containing MVBs travel on actin tracks. Future strategies targeting this pathway could provide further insight into exosome biology.

## Discussion

Exosomes have recently come forth as important mediator of cellular communication governing progression of various inflammatory disorders, and their increasing relevance to human pathologies command better tools for understanding their biology as well as for therapeutic purposes. Here, we provide first evidence that cholesterol lowering drug simvastatin could inhibit exosome synthesis and trafficking. Our results also suggest that some well-known anti-inflammatory and atherosclerosis preventive effects of statins may be linked to inhibition of exosome secretion.

We had previously reported that exosomes actively play a proinflammatory role in asthma pathogenesis and speculated that molecules capable of reducing exosome secretion might play a protective role in asthma. ^10^ In this study, we report that simvastatin mediated exosome reduction indeed result in protective phenotype in murine model of asthmatic airway inflammation, which is also supported by recent reports of beneficial role of statins in human subjects ^26^ and other experimental studies that focused on nitric oxide metabolism. ^22^ However, such *in vivo* models are complex and it is difficult to know whether the reduction in exosome secretion led to reduction in inflammation or *vice versa*. In support of an exosome-mediated effect, we found that the culture supernatant from simvastatin treated monocytes was diminished in exosomes and pro-inflammatory exosomal miRNA content, and also lacked the ability to induce endothelial migration.

While we found a number of interesting leads, the mechanisms by which statins potentially inhibit exosome secretion is not completely clear. Our finding that simvastatin regulated multiple proteins of exosome production machinery suggests the existence of a dedicated inter-connected protein network for exosomal production, managed by few key master regulators. While we started the study in the belief that inhibiting cholesterol synthesis may attenuate exosomal membrane formation, this seems too simplistic. Mevalonate supplementation was unable to restore exosome secretion in mouse lungs **(Figure 4A)**. Clearly, these data do not exclude the possibility that simvastatin may exert other functional effects through alternative pathways as well. We also understand that full potential of such discoveries can be exploited only in conjunction with development of tools for their selective targeting as well.

Our finding that exosome containing MVBs may travel on actin networks and simvastatin treatment significantly alters the length of these linear structures and their membrane association together offers exciting new directions and tool to look for novel proteins regulating exosomes via altering MVB movements.

In summary, this study identifies simvastatin as a potential tool to target pathway of exosomes, and significantly extends the role of simvastatin than just being a cholesterol lowering drug to a potential adjuvant for exosome dominated pathologies.

## Materials and Methods

### Cell Lines

Bronchial Epithelial cell line BEAS-2B and human epithelial carcinoma cell-line NCI H-1299 was procured from ATCC (Middlesex, UK). BEAS-2B was cultured in BEGM media supplemented with bullet kit from Lonza, THP-1 and H-1299 cells were maintained in RPMI 1640 supplemented with 10% FCS. HUVECs were isolated from human umbilical cord and were cultured in M199 media supplemented with ECGF (Sigma, USA). Experiments with Human umbilical cords were performed as per guidelines and protocols approved by the Institutional Human Ethics Committee.

### Animals

Male BALB/c mice (8-10 weeks old) were obtained from National Institute of Nutrition (Hyderabad, India) and acclimatized for a week prior to the experiments. All animals were maintained as per guidelines and protocols approved by the Institutional Animal Ethics Committee.

### Antibodies

CD-63, Alix, Tsg-101, Hsp-70, β-actin and GAPDH were purchased from Santacruz Biotechnology (Santa Cruz, CA). Fluorescently labeled Annexin-V, CD-81 (FITC), CD-63 (FITC), CD-63 (PE) and CD-9 (FITC) were purchased from BD biosciences (San Jose, CA).

### Development of OVA-sensitized Mouse Model of asthma and treatment of mice with simvastatin

Mice were sensitized and challenged as described earlier ^28^ (**Figure S5 in the online supplement**). Mice were divided into four groups as indicated, each group (n=6) was named according to sensitization/challenge/treatment: SHAM/PBS/VEH (normal controls, VEH-vehicle) called ‘SHAM’, OVA/OVA/VEH (allergic controls, OVA, chicken egg ovalbumin, Grade V, Sigma, USA) called ‘OVA’, OVA/OVA/Simvastatin (allergic mice treated with 40 mg/kg/dose Simvastatin, Sigma, USA) called ‘Statin’ and OVA/OVA/Simvastatin + Mevalonate (allergic mice treated with 40 mg/kg/dose Simvastatin and 20 mg/kg/dose mevalonate Sigma, USA) called ‘Mevalonate’ respectively.

### Exosome Isolation

Exosomes were isolated using a series of centrifugation and ultracentrifugation techniques as described elsewhere ^26^ with the modification wherein the supernatant from 10,000g fraction was filtered with a 0.2μm membrane before subjecting it to ultracentrifugation at 100,000g for 2 hours. The pellet was then washed with a large amount of PBS and then resuspended in 200μl of PBS, which was then sucrose density gradient purified. The exosome pellet was suspended in lamelli buffer when used for western blotting.

### Semi-quantitative detection of exosomes by bead-based assay

For semi-quantitative detection of exosomes, antibody coated beads were used as described earlier. ^14^ Briefly, 20,000 anti-CD-63 antibody coated beads were washed in 2% BSA and then incubated with 10,000g supernatant of BALF or culture supernatant overnight. Next day, the bead bound exosomes were detected using surface proteins for exosomes or phosphatidylserine on their surface, using flow cytometer.

### Flow cytometry

For surface labeling, cells were incubated with the antibodies diluted 1:25 in staining buffer for 30 minutes on ice, followed by a PBS wash, after which the cells were fixed with 2% paraformaldehyde. For intracellular labeling, cells were fixed and permeabilized, followed by staining. For total (surface+intracellular) CD-63 staining, initially the surface labeling was carried out as mentioned above. After the antibody incubation, cells were fixed and permeabilized and then the protocol for intracellular labeling was carried out.

### Western blot

Total cell protein was extracted and was resolved onto a polyacrylamide gel, which was then transferred onto a nitrocellulose membrane. The membrane was blocked with 5% skimmed milk and then probed for proteins of interest.

### Measurement of Airway hyper-responsiveness (AHR)

AHR in the form of airway resistance was estimated in anesthetized mice using the FlexiVent system (Scireq, Canada) which uses a computer-controlled mouse ventilator and integrates with respiratory mechanics as described previously. ^25^ Final results were expressed as airway resistance with increasing concentrations of methacholine.

### Lung Histology

Formalin-fixed, paraffin-embedded lung tissue sections were examined for airway inflammation, goblet cell metaplasia and sub-epithelial fibrosis with Hematoxylin & Eosin (H&E), Periodic acid-Schiff (PAS) and Masson-Trichrome (MT) staining respectively as described previously. ^28^ Briefly, grades of zero to four were given for no inflammation (zero), occasional cuffing with inflammatory cells (one), when most bronchi or vessels were surrounded by a thin layer (1–2 cells) (two), a moderate layer (3–5 cells) (three), and a thick layer (more than 5 cells deep) (four) of inflammatory cells and an increment of 0.5 was given if the inflammation fell between two grades and total inflammation score was calculated by addition of both peribronchial and perivascular inflammation scores.

### Measurements of cytokines in lung homogenate

Lung homogenates were used for ELISA of IL-4, IL-5, IFN-γ, IL-10 (BD Pharmingen, San Diego, CA) and IL-13 (R&D systems, Minneapolis, MN) as per the manufacturer’s protocol. Results were expressed in picograms and normalized by protein concentrations.

### Immunohistochemistry

Paraffin embedded tissue sections were used for preparation of 5μm tissue slides and immunohistochemistry was performed as described in. ^10^ CD-63 antibody was used at a dilution of 1:100.

### Immunofluorescence

Cells were seeded onto 0.17mm coverslips and immunofluorescence was performed as described, ^28^ CD-63 (FITC-conjugated) was used at a dilution of 1:250.

For exosome uptake assay by HUVECs, exosomes isolated from THP-1 cells were labeled with DiO-C16 for 1 hour and then unlabeled dye was removed by washing with PBS. Purified labeled exosomes were isolated by floating DIO labeled exosomes on sucrose density gradient. THP-1 exosomes thus isolated were resuspended in M-199 medium and incubated with cultured HUVEC cells. After incubation for 4 hours, HUVEC cells were washed, fixed, and observed under confocal microscopy.

### Quantitative polymerase chain reaction protocol

Real Time PCR for microRNAs were performed with sybr-green using custom primers. Equal concentration of starting RNA was used from each treatment for measuring micro-RNA in supernatant, while micro-RNA in cells were normalized to 18s rRNA as internal control. Primer sequences used were mir-150 RT primer 5’-CTCAACTGGTGTCGTGGAGTCGGCAATTCAGTTGAGCACTGGTA-3’, mir-150 forward primer, 5’-ACACTCCAGCTGGGTCTCCCAACCCTTGTA-3’, mir-16 RT primer 5’-CTCAACTGGTGTCGTGGAGTCGGCAATTCAGTTGAGCGCCAAT A-3’, mir-16 forward primer, 5’-ACACTCCAGCTGGGTAGCAGCACGTAAATA-3’, mir-181 RT primer, 5’-CTCAACTGGTGTCGTGGAGTCGGCAATTCAGTTGAGACCCACCG-3’, mir-181 forward primer 5’-ACACTCCAGCTGGGAACATTCATTGCTGTCG-3’, universal reverse primer 5’-GTGTCGTGGAGTCGGCAATTC-3’.

### HUVECs transmigration assay

The migration ability of HUVEC was tested in a Transwell Boyden Chamber (6.5 mm, Costar). The polycarbonate membranes (8 μm pore size) on the bottom of the upper compartment of the Transwells were coated with 1% gelatin matrix. Cells were suspended in serum-free M-199 culture medium at a concentration of 4 × 10^5^ cells/ml, treated with or without simvastatin treated THP-1 MVs for 2 hr and then added to the upper chamber (4 × 104 cells/well). Simultaneously, 0.5 ml of M-199 with 10% FBS was added to the lower compartment, and the Transwell-containing plates were incubated for 4 hr in a 5% CO2 atmosphere saturated with H2O. At the end of the incubation, cells that had entered the lower surface of the filter membrane were fixed with 90% ethanol for 15 min at room temperature, washed three times with distilled water, and stained with 0.1% crystal violet in 0.1 M borate and 2% ethanol for 15 min at room temperature. Cells remaining on the upper surface of the filter membrane (nonmigrant) were scraped off gently with a cotton swab. Images of migrant cells were captured by a photomicroscope. Cell migration was quantified by blind counting of the migrated cells on the lower surface of the membrane, with five fields per chamber.

### Statistical analysis

Data are expressed as mean ± standard error (SE). Significance of differences between groups was estimated using unpaired Student t-test for two groups or ANOVA with post hoc testing and Bonferroni correction for multiple group comparisons. Statistical significance was set at p ≤ 0.05.

## ACKNOWLEDGEMENTS

Authors (AK, KK) acknowledge the help of CSIR for their fellowships. This work is supported by grants from Council of Scientific and Industrial Research (CSIR), India (Task Force Project BSC0116, MLP5502), Department of Science & Technology (GAP84) and DST-INSPIRE grant to AK. We also acknowledge St. Stephen’s Hospital, New Delhi for providing Human Umbilical Cords for HUVEC isolation.

## Author contributions

AK and BG conceptualized and established the hypotheses. AK designed the study, executed the experiments, performed data acquisition, analysis and interpretation, drafted the manuscript,
critically revised the manuscript and performed statistical analysis; SS and KK was involved in the study design, experiments and co-analysis of data; assisted critical revision of the manuscript and provided technical support; AA and BG were involved in conception and design of the study, interpretation of data, drafting of the manuscript, critical revision of the manuscript for important intellectual content, obtaining funding and supervision.

## Additional information

Author(s) declare no Competing financial interests.

## Figure Legends

**Figure 1. Simvastatin reduces exosomes secretion. (A-B)**, Cells at a concentration of 2x10^6^/well of a 6-well plate were treated with indicated concentrations of simvastatin in 2 ml of media for a period of 24 hours, after which the culture supernatant was harvested and 1ml from it was used for measuring exosomes. Secreted exosome levels in culture supernatant from simvastatin treated epithelial cells **(A)** and THP-1 monocytes **(B)**, measured as in **Figure S4 in the online supplement**. **(C)**, Levels of exosome associated Alix, Tsg-101 and β-actin in pelleted exosome fraction from supernatant of 10^7^ simvastatin treated cells. **(D)** Effect of simvastatin treatment on exosome associated CD9/CD81 and Annexin V in cell culture supernatant from IL13 (25 ng/ml) and simvastatin treated epithelial cells. Data in **A, B** and **D** represent the mean±SE from three independent experiments. Data in **C** is representative image from one of the two independent experiments. (∗p<0.05 vs Control and ¥ p< .05 vs rIL13). Sim: Simvastatin.

**Figure 2. Simvastatin directly alters the level of various exosome associated proteins. (A)** Western blots for Alix and CD63 levels in total cell protein with different doses of simvastatin. **(B)** Cell surface levels of CD63 and CD81 were measured by flow cytometry after treatment with various doses of simvastatin, E-cadherin was used as control surface marker. **(C)** Immunocytochemistry for CD63 on cells treated with indicated concentration of simvastatin. Sim: Simvastatin.

**Figure 3. Effect of simvastatin and mevalonate cotreatment on inflammatory parameters. (A)** Secreted exosome levels in BAL supernatant of mice from indicated groups. **(B-C)** Lung sections stained with hematoxylin and eosin **(H&E, B)** showing leukocyte infiltration, periodic acid–Schiff **(PAS, C)** for collagen deposition. **(D)** Airway resistance with increasing concentrations of methacholine 12h after the last challenge. **(E-F)** Effect of indicated treatments on total leukocyte count **(E)** and differential leukocyte count enumerated by morphological criteria **(F)**. **(G)** Ova specific serum IgE levels measured by ELISA. **(H)** Cytokines IL-13 and IL-4 measured in pulmonary homogenate. Stains in **(B, C)** shown at 20X magnification. **Br**, Bronchus. Results **(A, D, E, F, G, H)** are the mean ± SE for each group from two experiments with 4-6 mice in each group, (**^∗^**, p<0.05 vs SHAM and ¥ p<0.05 vs OVA.), Sim: Simvastatin (40 μmg/kg/dose), Mev: Mevalonate (20 mg/kg/dose).

**Figure 4. Simvastatin reduces exosome production from monocytes and attenuates exosome-enclosed mir-150 mediated endothelial cell migration**. 1x10^6^ of THP-1 cells were seeded and treated with 0.3μM of simvastatin and 2μM GW4869 for a period of 24 hours. Cell pellet and supernatant was harvested and used separately for RNA isolation. Presence of indicated micro-RNAs was determined using qRT-PCR. Simvastatin mediated reduction of exosomes secretion from THP-1 monocytes results in lower levels of secretory miRNAs **(A)** but not intracellular miRNAs **(B). (C)** Uptake of DIO-labeled THP-1 derived exosomes (10 μg/mL) by HUVECs. **(D)** Relative mir-150 levels in HUVECs with indicated treatment **(E),** Simvastatin mediated reduction in exosome secretion by THP-1 monocytes results in lower mir-150 levels in HUVECs incubated with exosomes from simvastatin treated THP-1 in comparison to exosomes from same number of untreated THP-1 control cells, and consequent reduction in migration of endothelial cells. (∗p<0.05 vs Control in **A**, **D** and **E2.** ¥p<0.05 vs Ctrl+MV in **D** and **E2**). Sim: Simvastatin, **Control:** Control HUVEC.

**Figure 5. Simvastatin alters localization of CD63-positive compartments in cells.** Representative Immunohistochemistry images for CD63 from lung tissue sections of OVA and OVA/Simvastatin treated mice in **(A)** epithelial cells and macrophages **(B)**. **(C),** Representative confocal images of CD-63 levels and localization post treatment with simvastatin. Arrows indicates CD63 localization pattern. **(D)** Representative images showing CD63-EGFP distribution and its association with linear beeline like structures **(D1-D2, inset)** in control and simvastatin treated cells in subplasmalemmal region, detected by TIRF microscopy and quantification of relative length **(D3). (E)** Co-localization of CD63 with actin nucleation sites as visualized by confocal microscopy. **(F)** Localization of CD63 with actin in absence and presence of actin polymerization enhancer Jasplakinolide (100 nM). Images in **(E, F)** shown at 63X while **(A, B, C, D)** shown at 100 X magnification. (∗p<0.05 vs Control in **Figure D3**). Sim: Simvastatin.

